# Distinct neural mechanisms construct classical versus extraclassical inhibitory surrounds in an inhibitory nucleus in the midbrain attention network

**DOI:** 10.1101/2020.03.13.990952

**Authors:** Hannah M. Schryver, Jing Xuan Lim, Shreesh P. Mysore

## Abstract

Inhibitory neurons in the midbrain spatial attention network, called isthmi pars magnocellularis (Imc), control stimulus selection by the sensorimotor and attentional hub, the optic tectum (OT). Here, we investigate in the barn owl how classical as well as extraclassical (global) inhibitory surrounds of Imc receptive fields (RFs), fundamental units of Imc computational function, are constructed. We find that focal, reversible blockade of GABAergic input onto Imc neurons disconnects their extraclassical inhibitory surrounds, but, surprisingly, leaves intact their classical surrounds. Subsequently, with paired recordings and iontophoresis, first at spatially aligned site-pairs in Imc and OT, and then, at mutually distant site-pairs within Imc, we demonstrate that classical inhibitory surrounds of Imc RFs are inherited from OT, but their extraclassical inhibitory surrounds are constructed within Imc. These results reveal key design principles of the midbrain spatial attention circuit, and attest to the critical importance of competitive interactions within Imc for its operation.

## INTRODUCTION

Animals behave in complex environments and are constantly faced with multiple competing stimuli. Selecting the location with the most “important” or highest priority stimulus to guide behavior at any instant is an essential part of adaptive behavior, and operates upon the foundation of a spatial map of relative stimulus priority ^1-4^. Equally essential is the processing and representation of the stimulus at the selected location. A core characteristic of neurons involved in spatial selection that impacts both these functions is their spatial receptive field (RF), defined as the subset of the spatial locations that a neuron responds to selectively. The excitatory center and classical inhibitory surround of the RF together control the responses of neurons to a stimulus inside the RF, whereas the extraclassical surround controls the modulation of the neuron’s responses by a competing stimulus outside the RF. Thus, understanding how classical and extraclassical surrounds are constructed is essential for understanding how neurons involved in selection achieve both competitive selection among multiple competing stimuli as well as the processing and representation of the selected target. The optic tectum (OT, or superior colliculus, SC, in mammals) is a major sensorimotor hub in the midbrain (Fig. 1A). SC/OT neurons encode space topographically, and are also known to encode a spatial map of relative stimulus priority ^2,5-7^. The SC/OT is required for the control of spatial selection and selective attention when a target is present amidst distracters^8-10^, with OT neurons signaling the highest priority stimulus among competing stimuli categorically ^11-13^. Notably, these competitive interactions within the OT are controlled by long-range inhibition generated by GABAergic neurons in the nearby isthmi pars magnocellularis (Imc; Fig. 1AB ^14^. Specifically, focal inactivation of Imc neurons abolishes stimulus competition within the OT ^15,16^. Additionally, recent results demonstrate that the signaling of the strongest stimulus by the Imc occurs earlier, and is more categorical, than in the OT, further highlighting the importance of Imc to the function of the midbrain selection network.

**Figure. 1.**
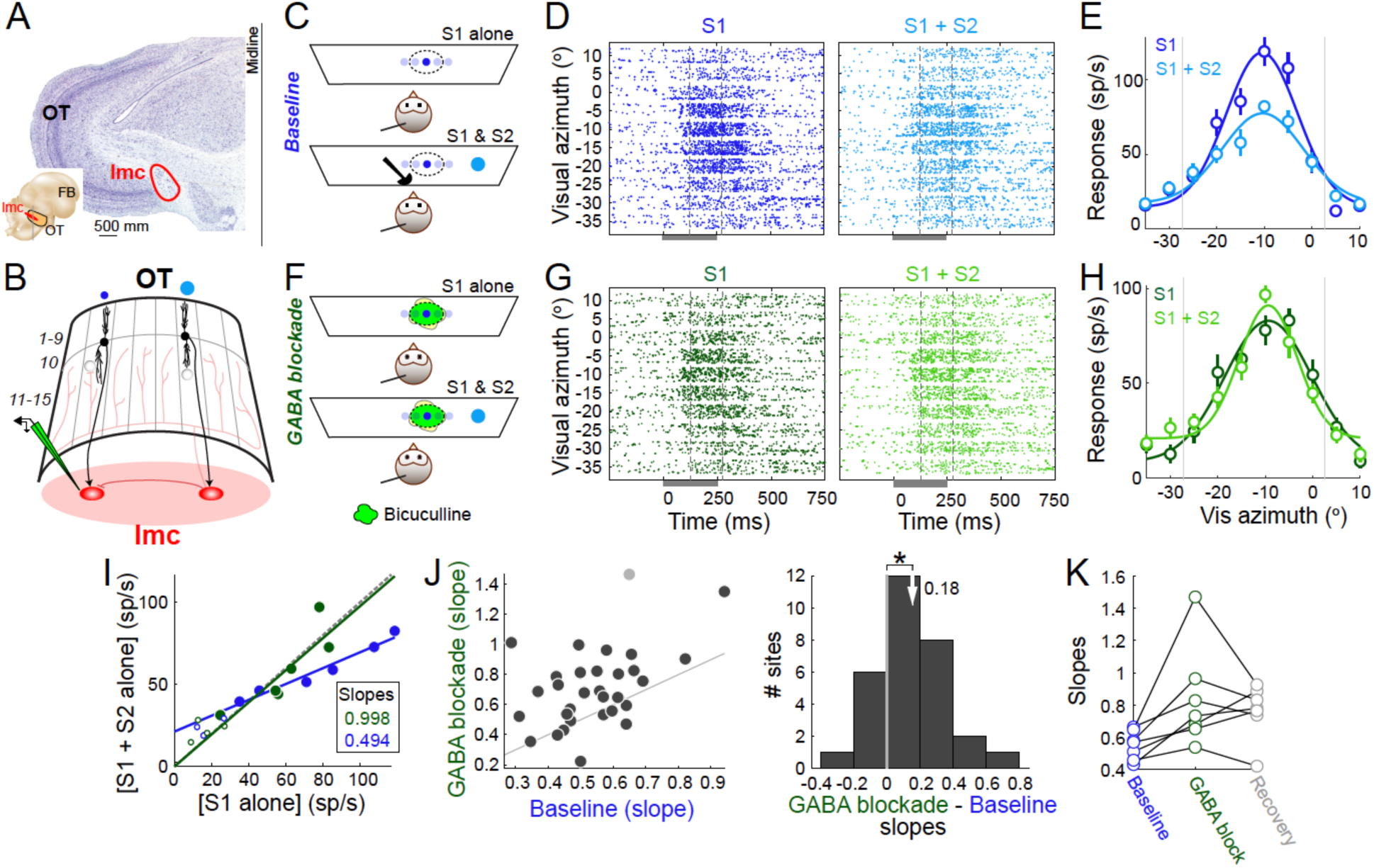
Imc’s extraclassical inhibitory surrounds are computed locally in the Imc. **(A)**Inset: Side view of schematic of barn owl brain. OT: optic tectum, Imc: nucleus isthmi pars magnocellularis, FB: forebrain. Gray vertical line: Coronal section through midbrain. <u>Main</u>: Nissl-stained coronal section of barn owl midbrain. “C”-shaped staining: layers of OT. **(B)**Schematic showing Imc-OT network connectivity and location of recording electrode. Curved sheet: OT; numbers correspond to layers (1-15). Columns: portions of OT (across layers) encoding adjacent spatial locations. Black neurons: OT layer 10 (OT_10_) neurons encoding distant azimuthal locations. Blue dots: stimuli presented at those two locations. Red ovals: Imc neurons. Translucent red projections: Projection pattern of the Imc neuron on the right to the OT; projections broadly innervate the OT (intermediate/deep layers of OT – 11 to 15, OT_id_), sparing only the portion (columns) of OT providing input ^14,25^. Arrows with sharp heads: excitatory projections; arrows with flat heads: inhibitory projections. Electrode symbol shows multi-barrel glass iontophoretic electrode, with bicuculline methiodide (green). **(C)**Stimulus protocol for measuring tuning curves in Imc without and with a distant competitor in the baseline condition. <u>Top</u>: Schematic showing owls viewing a tangent monitor. Angled line: recording electrode. Dotted oval: Spatial receptive field (RF) of recorded site. Dark blue dot: visual looming stimulus (S1), at various azimuthal locations (transparent blue dots). <u>Bottom</u>: Light blue dot: competing visual looming stimulus (S2) at a distant azimuthal location. Difference between dot sizes denotes differences in loom speed (strength) of stimuli; S2 always stronger than S1 (Methods). **(D**,**E)** Responses of an example Imc site in baseline condition. (D) <u>Left</u>: Raster plot of neural responses of an example Imc site to S1 presented at different azimuthal locations (tuning curve; C, top); loom speed of S1 = 6 °/s; S1 elevation = 5°). Negative values of azimuth indicate contralateral locations. <u>Right</u>: Raster plot of neural responses to S1 when S2 is also presented simultaneously (Fig. 1C, bottom; S2 loom speed = 10 °/s; S2 azimuth = −40 °, elevation= 5 °). Dark gray bar: Stimulus duration (250 ms). Dotted vertical lines: window of responses used for estimating response firing rates (125-275 ms). (E) Response firing rates of example site in E. Dark blue: S1 alone. Light blue: S1 and S2 presented simultaneously. Data represent mean +/- s.e.m. Solid lines: gaussian fits. Grey vertical lines: RF boundaries determined from S1 alone tuning curve (dark blue curve). **(F)** Stimulus protocol for measuring tuning curves in Imc without and with a distant competitor, during GABA blockade with bicuculline iontophoresis. Conventions as in C. Green blob: represents the focal nature of bicuculline iontophoresis. **(G**,**H)** Responses of same example Imc site in D,E, during GABA blockade. Dark green: S1 alone. Light green: S1 + S2. Other conventions as in D,E. **(I)**Scatter plots of responses of example Imc site to S1 alone versus to the paired presentation of S1 and S2. Blue data: baseline condition. Green data: GABA blockade condition. Large, filled dots: locations within the RF of the site. Small, open dots: locations outside the RF of the site. Straight lines: linear fits to responses for S1 locations within the RF; slopes are shown, and they indicate the extent of suppression (smaller slope – greater suppression). **(J)**Population summary. <u>Left</u>: Scatter plot of slopes in baseline versus GABA blockade conditions; n=30 (Gray dots – outliers; Methods). <u>Right</u>: Histogram of differences between slopes in GABA blockade and baseline conditions for each site (outliers removed). ‘*’: statistically significant; t-test comparing the mean (white arrow) to zero (gray line), p=0.0004. **(K)**Recovery. Summary plot of the slopes at a subset of Imc sites (n=7), measured in the baseline, GABA blockade, and recovery conditions.

In turn, Imc neurons, which have well-defined spatial receptive fields ^17,18^, exhibit both classical and extraclassical competitive surrounds ^19,20^. The responses of Imc neurons to two stimuli inside the RF are sub-additive and represent the average of their individual responses ^19^, indicating the presence of a classical inhibitory surround ^19,21-23^. In parallel, responses of Imc neurons to a stimulus inside the RF are divisively suppressed by a second stimulus anywhere outside the RF, demonstrating a global, extraclassical inhibitory surround. Whereas the source of the excitatory drive for Imc receptive fields are known to be neurons in layer 10 of the OT (OT_10_; ^24^, how the inhibitory surrounds of Imc receptive fields are constructed is poorly understood. Addressing this question is key to understanding how the map of relative stimulus priority and selection in the OT are orchestrated.

Here, we systematically dissect the mechanisms by which the inhibitory surrounds of Imc neurons in the barn owl are constructed. We do so in a series of experiments involving extracellular recordings in the Imc coupled with iontophoretic silencing of GABAergic input onto Imc neurons, of GABAergic input onto spatially aligned OT sites, or of excitatory (glutamatergic) input onto other/distant Imc sites. First, with iontophoretic blockade of GABAergic input on Imc neurons, we show that whereas global competitive surrounds of Imc neurons are abolished, surprisingly, the classical inhibitory surrounds are not affected. Then, with paired recording experiments at spatially aligned sites in the Imc and OT, we demonstrate that inhibition onto OT_10_ neurons – the sole source of excitatory input to Imc neurons – controls classical inhibitory surrounds of Imc RFs. Finally, with paired recording experiments at sites encoding for distant locations within the Imc, we demonstrate that distant Imc neurons control the global competitive surrounds in the Imc. Our results reveal that distinct mechanisms are involved in the construction of the inhibitory surrounds of Imc neurons: classical, local inhibitory surrounds are conferred onto Imc neurons by OT_10_ neurons, while global, competitive surrounds are constructed within the Imc using inhibition from distant Imc neurons.

## RESULTS

### Imc’s extraclassical inhibitory surrounds are computed locally in the Imc

We first investigated mechanisms underlying the extraclassical inhibitory surrounds exhibited by Imc neurons. Specifically, we asked if the previously reported reduction in Imc responses to a stimulus (S1) inside the RF, by a distant competitor (S2) presented outside the RF ^19,20^, was due to a comparison occurring at the Imc site itself, or if the response reduction reflected computations occurring elsewhere.

To this end, we conducted extracellular recordings in the Imc using multibarrel glass iontophoresis electrodes filled with a bicuculline methiodide solution (Fig. 1B, Methods). We recorded tuning curve responses at Imc sites using a single stimulus (S1), or while simultaneously also presenting a second, stronger stimulus far outside a site’s RF (S2; ∼30° away; Fig. 1C). Both S1 and S2 were visual looming stimuli; their loom speeds represent their strengths (Methods; S1 strength: 6 °/s, S2 strength 10 °/s). Trials with S1 alone, or S1 and S2 presented simultaneously, were interleaved randomly. Consistent with previous findings, responses to paired S1 and S2 presentations were significantly reduced compared to responses to S1 alone (Fig. 1D,E,I; slope=0.49, CI: [0.35, 0.637]). We then repeated these measurements following iontophoresis of bicuculline at the Imc recording site (Fig 1F, Methods), thereby blocking GABAergic synaptic transmission onto the recorded Imc neurons. We found that this abolished the reduction of responses due to the competitor S2, thereby disconnecting this Imc site’s extraclassical surround (Fig. 1G,H,I; slope=0.998, CI:[0.18 1.82]).

Across a population of tested Imc sites (n=30), we found that bicuculline iontophoresis consistently and significantly weakened this competitor-dependent response reduction (Fig. 1JK; p=0.0004, t-test against 0, eta^2^ = 0.14). We verified that these results were specifically due to drug iontophoresis by observing, at a subset of Imc sites, responses following recovery from bicuculline iontophoresis (Methods); we found that competitor-dependent response reduction returned to strong, near-baseline levels (Fig. 1K).

These results demonstrate that the reduction of Imc responses by a distant competing stimulus is due to suppression occurring via GABAergic synapses on Imc neurons. Global inhibitory surrounds are thus constructed locally at the Imc neurons themselves.

### Imc’s classical inhibitory surrounds are not computed locally in the Imc

We next investigated mechanisms underlying the classical surrounds exhibited by Imc neurons. Specifically, we tested if inhibition impinging onto Imc neurons, a common source of the classical surrounds of RFs in cortical and sub-cortical brain areas, mediates the construction of classical surrounds in Imc as well.

To this end, we examined the spatial tuning curves recorded in the previous experiment during the presentation of S1 alone, and compared the half-max widths of the tuning curves before and during iontophoresis of bicuculline (Fig. 2A-C). Should inhibitory influences onto Imc neurons be involved in creating the classical surround, we would expect that blockade of inhibition would widen the tuning curve, increasing the size of the half-max width ^21,26^. Notably, however, at an example site, we found no effect of bicuculline iontophoresis on Imc tuning curves (Fig. 2BC, half-max widths = 24.3° (baseline), 23.1° (GABA blockade)).

**Figure 2.**
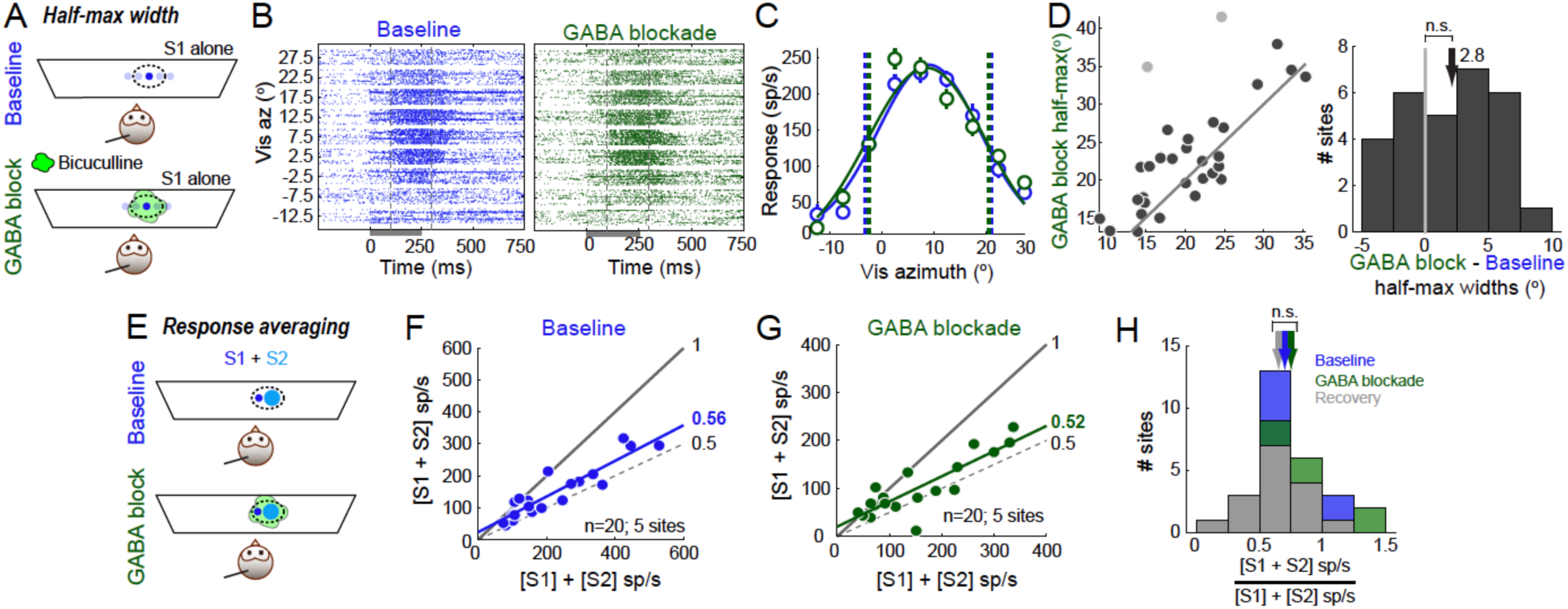
Imc’s classical inhibitory surrounds are not computed locally in the Imc. **(A-D)** Assessing the effect of GABA blockade on Imc classical inhibitory surrounds by using single stimulus, tuning curve measurements. (A)Stimulus protocol for measuring tuning curves in Imc in the baseline (Top) and GABA blockade (Bottom) conditions. Conventions as in Figure 1C. (B)Raster plots of neural responses of an example Imc site to S1 presented at different azimuthal locations (tuning curve; A, top) in the baseline (Left) and GABA blockade (Right) conditions. Positive values of azimuth indicate contralateral locations. (Site different from example site in Figure 1D,G.) Loom speed of S1 = 6 °/s; S1 elevation = 25 °. Conventions as in Figure 1D,G - left. (C)Response firing rates of example site in B. Blue: baseline; green: GABA blockade. Dotted vertical lines: Half-max widths, baseline = 24.3°. GABA blockade =23.1°. Other conventions as in Figure 1E,H. (D)Population summary. <u>Left</u>: Scatter plot of half-max widths of tuning curves in baseline versus GABA blockade conditions; n=29 (Gray dots – outliers; Methods). Note that these data are from the same sites as in Figure 1. <u>Right</u>: Histogram of differences between half-max widths in GABA blockade and baseline conditions for each site (outliers removed). ‘n.s.’: not significant; sign-test comparing the median (black arrow) to zero (gray line), p=0.132. **(E-H)** Assessing the effect of GABA blockade on Imc classical inhibitory surrounds by using a two-stimulus (response integration) protocol. (E)Stimulus protocol in which two stimuli (S1 and S2, filled dots) are presented simultaneously inside the spatial RF of the Imc site (dashed oval), in the baseline (Top) and GABA blockade (Bottom) conditions. Responses are compared to those obtained by presenting S1 and S2 individually (not shown). Other conventions as in Figure 1C. (F,G) Population summary (n=20 paired presentations). Scatter plot of responses to the presentation of S1 and S2 simultaneously, versus the sum of the responses to S1 and S2 presented individually. F – Baseline condition, G – GABA blockade condition. Black line: represents when the paired response equals the sum of the individual responses; slope =1. Dashed line: represents when the paired response equals the average of the individual responses; slope = 0.5. Colored lines: best fit to the data. Slopes of the best-fit lines are indicated. (H) Population summary. Histogram of ratios of the paired responses to the summed responses. Arrows: mean ratios. Blue: baseline; green: GABA blockade; grey: recovery. ‘n.s.’: not

Across the population of tested Imc sites (n=29), we found no systematic effect of bicuculline iontophoresis on the half-max widths of spatial tuning curves (Fig. 2D; p=0.14; sign-test against 0). Since these data were obtained at the same sites as in Figure 1, and in a randomly interleaved manner with that data, we ruled out the possibility that the drug (or the iontophoresis protocol) was ineffective at blocking GABA synapses. A potential remaining confound was that, perhaps, the half-max width was not a sufficiently sensitive metric to detect real changes in the classical inhibitory surround. To test this, we performed an additional experiment at a separate set of sites. We tested the effect of bicuculline iontophoresis on the response averaging previously reported when two stimuli are presented simultaneously inside the RF ^19^ (Methods). We found that Imc responses to two stimuli inside the RF were subadditive, a result dependent of the operation of the classical inhibitory surround, and specifically, represented an averaging of the responses to the stimuli presented individually (Fig. 2EF: slope of the scatter plot of the responses to S1 and S2 presented together against the sum of the responses to S1 and S2 presented individually =0.56, CI=[0.44,0.68]). Notably, bicuculline iontophoresis onto the Imc sites did not alter this response averaging effect (Fig. 2G: slope = 0.52, CI=[0.37,0.68]; Fig. 2H: ANOVA, F=0.72, p=0.4936, n=20).

These results together demonstrate that GABAergic inhibition onto Imc neurons does not participate in constructing their classical inhibitory surround.

### Imc’s classical inhibitory surrounds are inherited from the OT

Since in-situ GABA blockade did not alter the classical inhibitory surrounds of Imc neurons, we next investigated another source for the construction of these surrounds. Specifically, we asked if the classical surrounds are the result of modulation of the excitatory drive into Imc neurons (rather than inhibition onto them). To test this possibility, we selectively manipulated the excitatory input to Imc neurons while simultaneously recording their responses. Neurons in layer 10 of the OT (OT_10_) provide the only source of excitatory input to Imc neurons ^24^. Therefore, we made paired recordings in Imc and OT_10_, at sites that were spatially aligned (Methods; Fig. 3A, top). We then recorded spatial tuning curves (Fig. 3A, bottom) simultaneously in Imc and OT_10_, with and without iontophoresis of bicuculline onto OT_10_ neurons.

**Figure 3.**
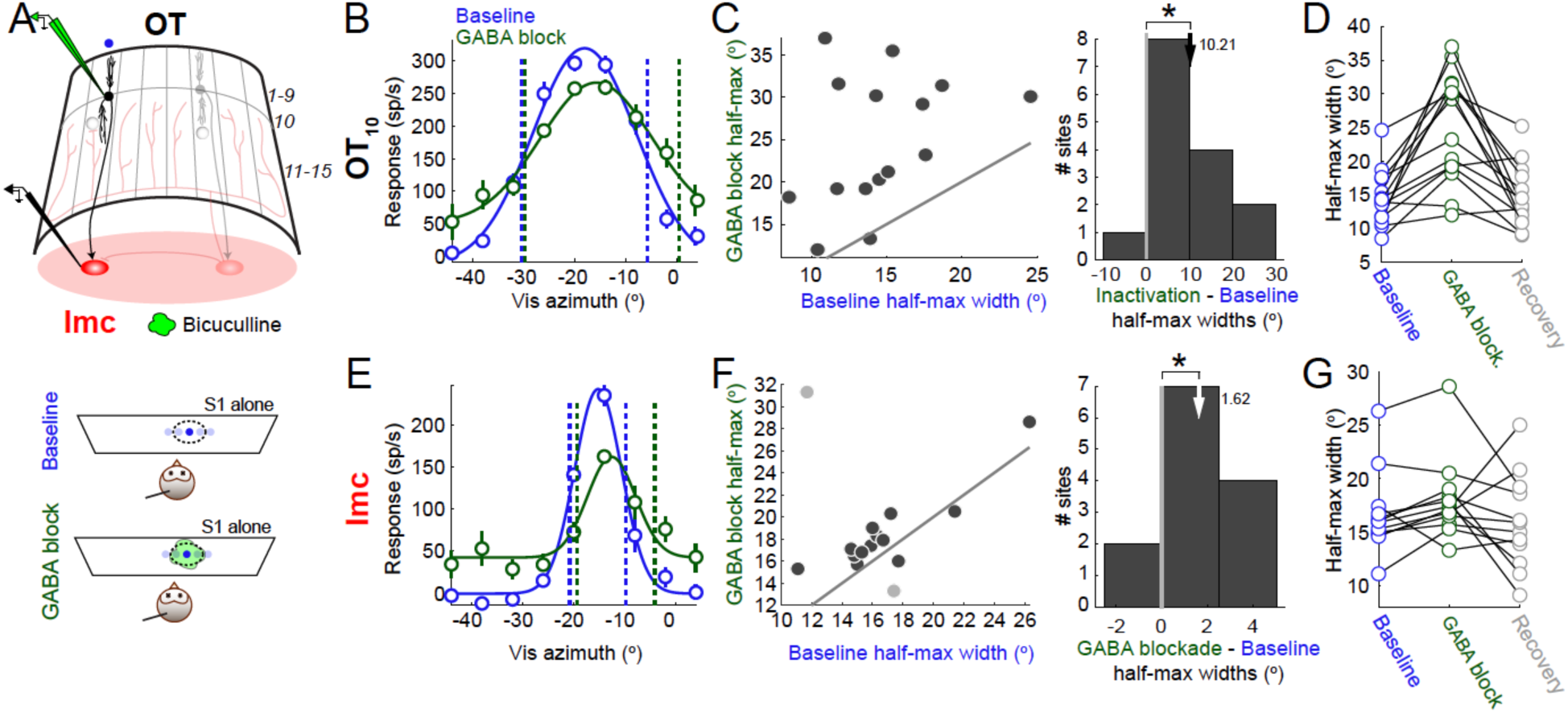
Imc’s classical inhibitory surrounds are inherited from OT_10_. **(A)**<u>Top:</u> Schematic showing Imc-OT network connectivity and location of recording electrodes. Electrode symbols show a multi-barrel glass iontophoretic electrode, with bicuculline methiodide (green) in OT_10_, and a second, tungsten electrode (in black) at a spatially aligned site in the Imc. Other conventions as in Figure 1B. <u>Bottom</u>: Stimulus protocol for measuring spatial tuning curves (simultaneously in spatially aligned Imc and OT_10_ sites) in the baseline condition (upper panel), and during GABA blockade in OT_10_ (lower panel). Conventions as in Figure 1C,F. **(B-D)** Responses measured at OT_10_ sites. (B)Firing rate responses of an example OT_10_ site to S1 presented at different azimuthal locations (tuning curve); S1 elevation = −24 ^º^; negative values of azimuth indicate contralateral locations. Blue: baseline; green: GABA blockade in OT_10_. Dotted vertical lines: OT_10_ half-max widths, baseline = 24.6°; OT_10_ GABA blockade =30.1°. Other conventions as in Figure 1E,H. (C)Population summary. <u>Left</u>: Scatter plot of half-max widths of OT_10_ tuning curves in baseline versus GABA blockade conditions; n=15. <u>Right</u>: Histogram of differences between half-max widths in OT_10_ GABA blockade and baseline conditions for each OT_10_ site. ‘*’: statistically significant; sign-test comparing the mean (black arrow) to zero (gray line), p=0.0001. (D)Recovery. Summary plot of the half-max widths at a subset of OT_10_ sites (n=14), measured in the baseline, OT_10_ GABA blockade, and recovery conditions. **(E-G)** Responses measured at Imc sites that were spatially aligned to the OT_10_ sites in B-D. All conventions as in B-D. (E) Imc half-max widths, baseline = 11.1° OT_10_ GABA blockade = 15.3°. (F) <u>Left</u>: Gray dots – outliers (Methods). <u>Right</u>: Histogram of differences between half-max widths in OT_10_ GABA blockade and baseline conditions for each Imc site (for data with outliers removed). ‘*’: statistically significant; t-test comparing the mean (black arrow) to zero (gray line), p=0.003. (G) Recovery. Summary plot of the half-max widths at a subset of Imc sites (n=12, outliers removed), measured in the baseline, OT_10_ GABA blockade, and recovery conditions.

We found that bicuculline iontophoresis onto OT_10_ neurons widened the OT_10_ spatial turning curve as quantified by an increase in the half-max width (Fig. 3B, example Imc site, half-max width = 11.1° (baseline), 15.3° (bicuculline iontophoresis)). This pattern held true across the population of tested OT_10_ sites (n=15), with bicuculline iontophoresis systematically increasing half-max widths of the spatial tuning curves (Fig. 3C, mean increase = 10.21°, p=0.0001, t-test against 0, eta^2^=0.4181). We confirmed that these results were due, specifically, to drug iontophoresis by observing that OT_10_ half-max widths returned to near-baseline values following recovery from bicuculline iontophoresis (Fig. 3D). (In addition, these results independently validated the efficacy of using half-max widths of spatial tuning curves as a means to assess the contribution of inhibitory inputs in shaping classical inhibitory surrounds.)

Next, we analyzed whether the spatial tuning curves measured simultaneously at the spatially aligned Imc sites were altered. Notably, we found a similar widening of Imc tuning curves by bicuculline iontophoresis onto OT_10_ neurons (Fig 3E, example Imc site paired with the OT_10_ site in Fig. 3B, half-max widths =24.6° (baseline), 30.1° (bicuculline iontophoresis)). Across the population of tested sites (n=15), we found a consistent widening of Imc sites when their paired OT_10_ sites underwent bicuculline iontophoresis (Fig. 3F-left, mean increase in Imc half-max widths = 2.44, p=0.035, sign-test against 0, eta^2^=0.08, with all the data points shown). To test if this result was driven entirely by the one data point representing a substantially larger increase in half-max width than all the other data points (Fig. 3F, left), we performed a standard outlier analysis (Methods; Fig. 3F-left, outlier datapoints in grey), and found a smaller but significant increase in half-max widths even after removing the outliers (Fig. 3F-right, mean increase in Imc half-max widths = 1.62°, p=0.0032, t-test against 0, eta^2^=0.05, with only the black data points in 3F-left). We also found that this increase in Imc tuning curve widths was the result, specifically, of drug iontophoresis (Fig. 3G).

Thus, silencing inhibition onto OT10 neurons affected the classical surrounds of their RFs, and modulated their output as a function of spatial location of stimulus S1. This, in turn, altered the excitatory input to Imc neurons and impacted their classical inhibitory surrounds. These results demonstrated that Imc classical surrounds are constructed via modulation of their excitatory drive, establishing inhibition onto OT_10_ (but not in Imc) as the primary source of Imc’s classical inhibitory surround.

### Imc’s extraclassical inhibitory surrounds are constructed using inhibition from other (distant) Imc neurons

Since in-situ GABA blockade interrupted the extraclassical inhibitory surrounds of Imc neurons, we were next interested in identifying the source of this inhibition. Past work in slice has shown the presence of long-range inhibitory projections between Imc neurons ^14,27^. To test if inhibition from other Imc neurons encoding for distant locations controls extraclassical inhibitory surrounds in Imc, we conducted paired recordings at two mutually distant sites within the Imc (Fig. 4A; Methods; sites > 20° apart). Using the same stimulus protocol as in Figure 1 (Fig. 1AB), we recorded tuning curve responses of neurons at one Imc site (site A) using stimulus S1, while simultaneously presenting S2 at a location encoded by a distant Imc site (site B). We then compared responses at Imc site A without and with iontophoretic silencing of Imc site B with kynurenic acid (Methods).

**Figure 4.**
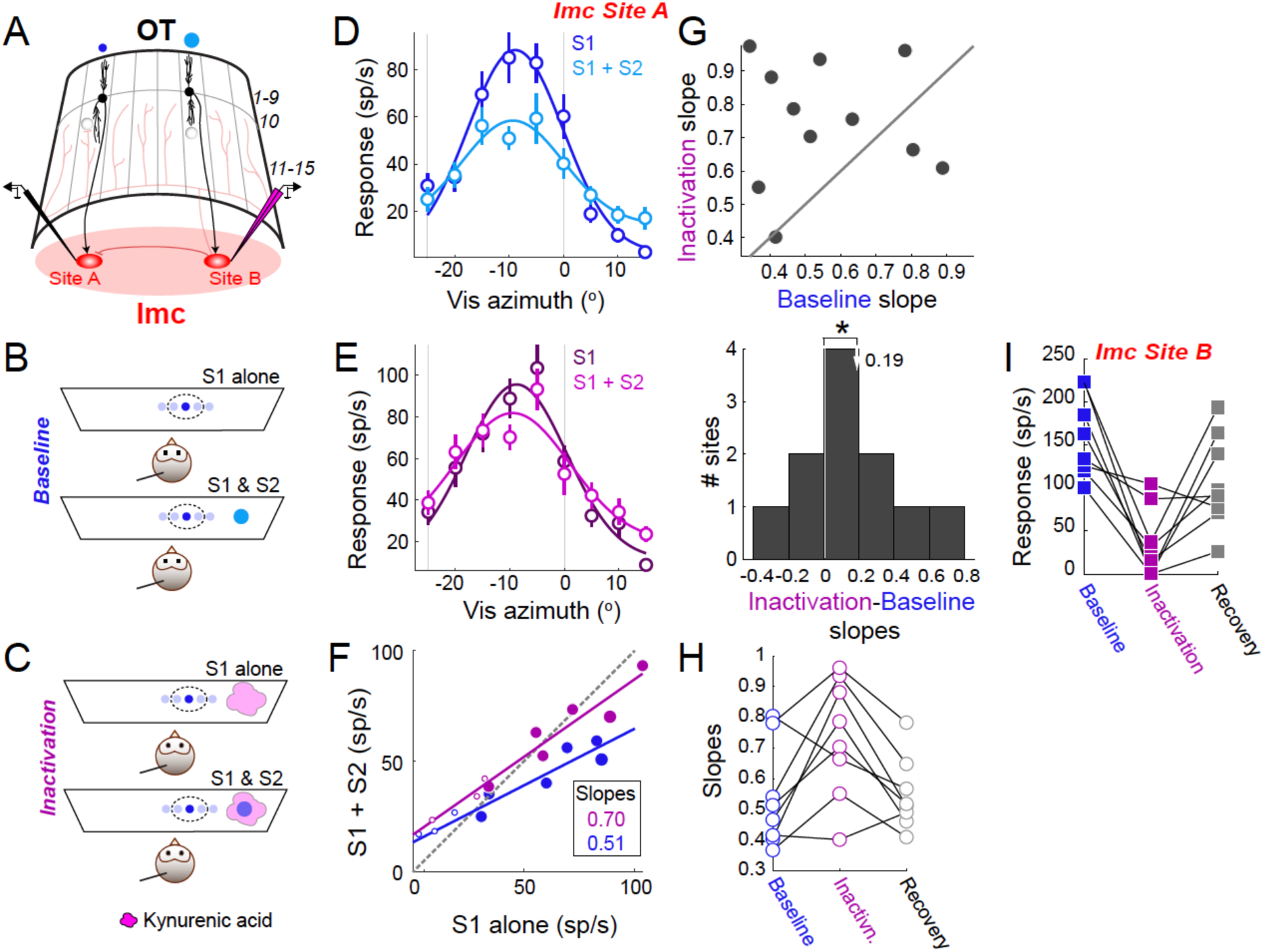
Imc’s extraclassical inhibitory surrounds are constructed using inhibition from other (distant) Imc neurons. **A)** Schematic showing Imc-OT network connectivity and location of recording electrodes. Electrode symbols show a tungsten electrode (in black) at Imc site A, and a multi-barrel glass iontophoretic electrode, with kynurenic acid (for silencing excitatory synaptic drive), at a distant Imc site B. Other conventions as in Figure 1B. **(B**,**C)** Stimulus protocol for measuring spatial tuning curves at Imc site A (using stimulus S1), without and with a distant competitor (S2) presented at a location encoded by Imc site B. (B) Baseline condition. (C) Imc site B inactivation condition. Conventions same as Fig. 1C. **(D-H)** Responses measured at Imc sites A. (D)Baseline condition. Firing rate responses of an example Imc site A to S1 presented at different azimuthal locations (tuning curve; S1 elevation = 30 °)e, without (dark blue) and with (light blue) the simultaneous presentation of a distant competitor S2 (azimuth = - 40°, elevation = 40°); negative values of azimuth indicate contralateral locations. Loom speeds: S1 = 6 °/sec; S2 =10 °/sec. Grey vertical lines: RF boundaries determined from S1 alone tuning curve (dark blue curve). Other conventions as in Figure 1E. (E)Firing rate responses of same example Imc site A as in D, but during inactivation of Imc site B. Dark magenta: tuning curve without S2; pink: tuning curve with S2. All other conventions as in D. (F)Scatter plots of responses of example Imc site A to S1 alone versus to the paired presentation of S1 and S2. Blue data: baseline condition. Magenta data: inactivation of Imc site B blockade condition. Straight lines: linear fits to responses for S1 locations within the RF; slopes are shown, and they indicate the extent of suppression (larger slope – weaker suppression). Other conventions as in Figure 1I. (G)Population summary. <u>Top</u>: Scatter plot of slopes at Imc sites A in baseline versus site B inactivation conditions; n=11. <u>Bottom</u>: Histogram of differences between site A slopes in the site B inactivation versus baseline conditions. ‘*’: statistically significant; t-test comparing the mean (white arrow) to zero (white line), p=0.04. (H)Recovery in site A. Summary plot of site A slopes at a subset of Imc sites (n=8), measured in the baseline, site B inactivation, and recovery conditions. **(I)**Inactivation of site B. Summary plot of site B responses to S2, measured in the baseline, site B inactivation, and recovery conditions (n=8).

We found that in the baseline condition, S2 effectively suppressed S1 tuning curve responses recorded at site A (Fig 4D,F, example site, slope=0.51, CI=[0.19,0.84]). However, focally inactivating Imc site B (encoding the competitor stimulus S2), abolished S2’s suppressive effect on responses of Imc site A (Fig. 4E,F; slope=0.70, CI=[0.33,1.1]). Across the population of tested site pairs (n=11 site A-site B pairs), we found consistently that iontophoresis of kynurenic acid at Imc site B significantly weakened the suppression of responses at Imc site A (Fig. 4G; mean slope increase of 0.19; p=0.04, t-test of inactivation vs. baseline, eta^2^=0.22). We found that these results were due, specifically, to drug iontophoresis, as evidenced by the return of S2-dependent suppression of responses at site A to near-baseline levels following recovery from kynurenic acid iontophoresis (Fig. 4H). Additionally, kynurenic was effective at inactivating Imc site B: responses were nearly abolished under kynurenic acid, and recovered following turning off of the drug (Fig. 4I).

Together, these results demonstrate that extraclassical inhibitory surrounds of Imc neurons are constructed (in the Imc) using long-range inhibition from distant Imc neurons, thereby establishing intra-Imc inhibition as the source of Imc’s extraclassical inhibitory surrounds.

## DISCUSSION

This study uncovers the mechanistic implementation of classical as well as global inhibitory surrounds of key inhibitory neurons in the midbrain spatial attention network, namely the Imc. These neurons form part of the tecto-fugal pathway for visuomotor (and more generally, sensorimotor) processing in vertebrates ^10^. In the parallel, thalamo-cortical pathway of visual processing, only recently was the debate resolved whether the classical surrounds of simple cells in V1 ^28^ are constructed by an appropriate organization of excitatory inputs from multiple upstream LGN cells, or instead, are constructed within V1 due to the action of local inhibitory neurons ^29^. Selective manipulation of inhibitory neurons within V1 in mice showed that cortical inhibition played a key role in the construction of the classical surround ^30^, thereby establishing a clear mechanism for a basic function of V1 neurons. In the tecto-fugal pathway, in addition to the SC/OT being involved in sensory processing, it plays a critical role in the competitive selection of the target for spatial attention. In turn, the Imc neurons control competitive interactions within the OT.

Recently, it was demonstrated that not only do Imc neurons supply inhibition to the OT, but also that this inhibitory output reflects computations occurring within the Imc, rather than being a sign-flipped version of excitatory inputs into Imc ^19,20^. Indeed, the Imc itself appears to encode a map of relative stimulus priority ^19,20^, much like the SC/OT – the first reported in inhibitory neurons, to the best of our knowledge. In this context, the construction of the responses of Imc neurons, and specifically, of their receptive fields is a critical question. Past work involving recordings in the Imc paired with focal iontophoretic silencing of OT_10_ neurons clearly identified OT_10_ as the source of Imc’s excitatory drive ^24^. However, although that study also examined the effect of iontophoretic application of bicuculline on Imc responses ^24^, those results are difficult to interpret in the context of the goals here. In that study, data obtained from the presentation of a single stimulus inside the RF, as well as from the presentation of two stimuli (one inside and one outside the RF) were combined together and reported as a single result. Classical versus global (extraclassical) surrounds have fundamentally different properties - in terms of their function, strength profiles and spatial scope, and requirements of the underlying circuitry. As a result, the properties of one cannot predict those of the other ^21^, highlighting that it is critical to consider separately results from single stimulus versus two-stimulus (competition) protocols, to disambiguate the mechanistic underpinnings of the respective inhibitory surrounds.

Doing so, here, revealed that stimulus competition is achieved locally in the Imc via inhibitory projections among Imc neurons. These long-range projections ^14,27^ create global inhibitory surrounds of Imc RFs. By contrast, our results from two different experimental tests revealed that no aspect of the classical inhibitory surround of Imc neurons is computed locally within the Imc – one test involved the comparison of the half-max widths of the single-stimulus tuning curves without and with blockade of synaptic inhibition, and the other involved comparison of (the sub-additivity of) Imc responses to two stimuli presented simultaneously within the RF. This finding is consistent with the observation in slice experiments that Imc neurons do not appear to receive projections from nearby Imc neurons ^27^. Instead, our results showed that the representation of a single stimulus by Imc neurons is purely a reflection of the excitatory input from OT_10_ to Imc, with the classical inhibitory surrounds of Imc RFs being inherited entirely via this excitatory input.

An interesting question that arises from these findings is why the Imc-OT circuit might be organized this way. In other words, why doesn’t the Imc inherit both classical and global inhibitory surrounds from the OT, or alternatively, why doesn’t it compute both locally? The fundamental function of the Imc, namely, orchestrating selection in the midbrain attention network, and specifically, in the OT, offers a plausible potential explanation. Imc is the sole source of competitive inhibition to the OTid and is necessary for the OTid to signal the highest priority stimulus ^15,16^. Additionally, Imc itself performs stimulus competition, does so earlier and more categorically than the OTid ^19,20^. Moreover, this competition within the Imc has been proposed, by modeling work, to be critical for mediating flexibility of selection boundaries in the OT ^31^. In other words, competition within the Imc appears to offer important advantages to signaling of the highest priority stimulus by the OTid. If true, having a dedicated circuit mechanism in the Imc that can implement extraclassical inhibitory surrounds and achieve stimulus competition locally, would appear to be beneficial. By contrast, in the case of classical inhibitory surrounds: Imc neurons have been shown to not send inhibition to the portion of the OT space map from which they receive input ^14,25^, a feature has recently been demonstrated to be required for the categorical nature of the signaling of the highest priority stimulus by the OTid ^25^. As a result of this anatomical feature, Imc does not participate in shaping the OTid’s responses to single stimuli. Considering this, not having a dedicated mechanism in the Imc for generating classical surrounds locally avoids ‘wasteful’ circuitry that would not sub-serve the core functional role of Imc. In other words, the computation of global surrounds, but inheritance of classical surrounds, may represent an efficient circuit implementation for stimulus selection in the midbrain attention network. This idea builds on the central premise that intra-Imc interactions are critical for stimulus selection by the OT, one that will be important to assess directly in future work with selective causal manipulations to the Imc-OT circuit.

## Acknowledgments

This work was supported in part by funding from NIH R01 EY027718. SPM and HMS designed the research, HMS and JXL performed experiments, HMS analyzed the data, SPM and HMS wrote the paper. The authors declare no competing interests.

## METHODS

### Experimental Design

The goal of this study was to determine the mechanisms underlying the construction of classical and extraclassical inhibitory surrounds of Imc neurons. This was done by measuring spatial tuning curves at Imc neurons using a visual stimulus (S1, presented at various azimuthal locations within and immediately outside the receptive field, RF), with or without a second visual stimulus present outside of the RF (S2). This was done in baseline conditions and while microiontophoretically applying bicuculline methiodide (Sigma-Aldrich), a GABA_A_ receptor antagonist locally at the recording site. This allowed us to compare the effect of GABaergic input on local surrounds (S1 tuning curves) and competitive surrounds (S1 tuning curves when S2 was also presented simultaneously). We also examined how the responses to two stimuli inside the RF are integrated by Imc neurons by presenting S1 alone within the RF, S2 alone within the RF at multiple distinct locations from that of S1, and also presenting S1 and S2 within the RF simultaneously. We then compared such responses obtained in the baseline condition and while microiontophoretically applying bicuculline methiodide to the recording site.

We also examined the role of OT_10_ neurons in the construction of classical inhibitory surrounds of Imc neurons by applying bicuculline methiodide at OT_10_ sites while simultaneously recording tuning curves at both the OT_10_ site and a paired Imc site encoding for the same area of sensory space (spatially aligned site). Finally, to examine the role of Imc neurons in mediating th extraclassical surrounds of other (distant) Imc neurons, we recorded from one Imc site encoding for S1 (site A) without and with S2 presented outside of the site’s RF. Here, a second, electrode was placed at the Imc site encoding for S2 (spatially misaligned site B). We then compared the responses of Imc sites A at baseline and when the paired sites B were inactivated using kynurenic acid.

### Neurophysiology

Six adult barn owls (Tyto alba; male and female) were used for electrophysiological recordings. All protocols and animal care were in accordance with NIH guidelines for care and use of laboratory animals and approved by the Johns Hopkins University Institutional Animal Care and Use Committee. Birds were shared across different studies and group housed in an aviary with a 12hr/12hr light/dark cycle. Before electrophysiological experiments, head bolts were mounted near the rear of the skull while owls were anesthetized with isoflurane (1-2%) and a mixture of nitrous oxide and oxygen (45:55). Birds were treated with an intramuscular injection of 0.1mL of Meloxicam and 0.1 mL of Butorphanol. All incisions were disinfected with Betadine and locally anesthetized with subcutaneously injected Bupivacaine. Bilateral craniotomies were performed and small plastic cylinders with removeable caps were placed on the skull to allow access to midbrain structures over multiple experiments. Polysporin antibiotic ointment was applied to any exposed brain surface and incisions. Owls were returned to the aviary following recovery from surgery and recovered for at least one week before experiments were performed.

On experiments days, owls were initially anesthetized with isoflurane (1-2%) and a mixture of nitrous oxide and oxygen (45:55) before being restrained in a flexible wrap. Birds were administered an intramuscular injection of 0.1mL of Meloxicam and 0.1 mL of Butorphanol. Birds were then secured in a sound-attenuating booth. Head-fixation was calibrated following published procedures ^32^ for each individual owl on experiment days. Owls were maintained on oxygen and nitrous oxide for the duration of the experiment, while isoflurane was turned off after the bird was secured. As recovery from isoflurane occurs well under 30 minutes after it is turned off, recordings were made in animals that were not anesthetized. When possible, nitrous oxide as well was turned off 5 minutes before data collection (it partitions out of blood rapidly, within a minute). Notably, previous work has demonstrated neural responses in the midbrain network do not differ under nitrous oxide tranquilization and non-tranquilized conditions ^13^.

The Imc is an oblong structure in the avian midbrain that is elongated along the rostrocaudal axis, parallel to the OT. Previous work has confirmed *in vivo* targeting of the Imc with fluorescent dye injection ^16^, and with electrolytic lesions ^18^, and established its location as approximately 500 µm medial to the medial-most part of the OT. Recording sites in the Imc were targeted by either navigating first to the optic tectum followed by the Imc using the OT’s topographic space map as reference ^19^, or by referencing reliable stereotaxic coordinates from prior experiments and verified on the basis of established distinct neural responses ^13,16,18,19,21^. For recording in OT_10_, OT layers were identified by distinctive neuronal responses published previously ^32^.

### Recordings

For Imc recordings, an electrode was positioned to enter the brain at a medial-leading angle of 5° to avoid a major blood vessel. During some paired Imc-Imc recordings, electrodes were also angled in the caudal-leading direction (2°-5°) to accommodate space for two electrodes. For paired OT_10_-Imc recordings, an electrode was positioned to enter the brain to record from OT_10_ was positioned at a 5° medial-leading angle to accommodate space for both OT_10_ and Imc electrodes. Extracellular activity was recorded primarily using multi-barreled glass electrodes with the central barrel containing a carbon fiber electrode for recording neural activity (Kation, Carbostar-3 Carbon fiber electrode). Paired recordings with two electrodes utilized one double-barreled glass electrodes to administer drug and record responses at one brain site, and one epoxy-coated tungsten microelectrode of high impedance (A-M Systems, 5 MΩ at 1kHz, 250 µm shaft diameter) to record response at the paired sites. Well-isolated single and multi-unit activity was recorded in the Imc. In a pilot experiments, we compared the results with data from multi-unit sites, versus with data from spike sorted single units, and found no differences (data not shown). For this reason, all data in this paper represent a combination of both single as well as mulit units. Spike times were recorded using Tucker-Davis hardware and analyzed using MATLAB.

### Microiontophoresis

Microiontophoresis was performed using a 1-channel iontophoresis box (DAGAN Corp PS-100) To achieve focal blockade of inhibitory synaptic input (in Imc and OT_10_), the GABA_A_ antagonist, bicuculline methiodide (Sigma, 10mM, 2.37-2.67 pH, mean pH = 2.54) was filled in one barrel of the multi-barreled electrode, and was microiontophoretically applied to the recording site. Bicuculline was ejected at currents ranging from 20 to 60nA (average 36.45 nA) at Imc sites or 80nA at OT sites for 15 minutes before data collection in the drug condition, and ceased for 25-35 minutes before data collection in the recovery condition. When not being ejected, bicuculline was retained at a current of −15nA. Responses were recorded using a carbon fiber electrode embedded in one of the barrels.

To achieve focal blockade of excitatory synaptic input (in Imc), the pan-glutamate receptor antagonist kynurenic acid (Sigma, 40mM, 8.5-9 pH) was filled in one barrel of the glass electrode and microointophoretically applied to the recording site. Kynurenic acid was ejected at −500nA for 15 minutes before before data collection in the drug condition, and ceased for 15 minutes before data collection in the recovery condition. When not being ejected, Kynurenic acid was retained at +15nA.

### Stimuli

Stimuli were black dots on a gray background presented on a 65” monitor. The stimuli expanded linearly over a duration of 250 ms to mimic looming (approaching) objects, and were used because previous work has established that looming dot stimuli evoke reliably strong responses in OT and Imc with relatively low response habituation ^13,21^. Stimuli were presented using MATLAB and Psychtoolbox (PTB-3; ^33,34^. The spatial locations of visual stimuli were defined by double pole coordinates relative to the midsagittal plane for azimuth or the visual plane for elevation ^32^.

To determine the extent of space encoded by a recorded site (in Imc or OT_10_), we collected two-dimensional RFs (azimuth x elevation) by presenting a single stimulus at various azimuthal and elevational locations. This stimulus was presented for 5 repetitions, with a duration of 250 ms and an inter stimulus interval of 1500 ms. Spatial locations at which a stimulus elicited higher firing rates compared to baseline were considered to constitute the site’s spatial RF and used for determining the placement of S1 (within the RF), S2 (inside or outside of the RF), and for determining the alignment of Imc and OT_10_ sites.

Tuning curves were measured by presenting S1 of a fixed strength (6 °/sec) at multiple azimuthal locations (mean = 9 locations) spanning the extent of a site’s RF. Individual locations at which S1 was presented were jittered by 2^º^ to reduce the effect of adaptation on neural responses. For examining extraclassical surrounds, S2 was presented outside the RF, at a distant location from S1 (mean distance from S1 = 32.8°). S2 loom speed (10 °/sec) was stronger than that of S1. Presentations of S1 alone, and S1 with S2, were interleaved pseudorandomly. Stimuli were presented for 15 repetitions for collecting the data in Figures 1-4. For examining classical surrounds using the two-stimulus response integration approach, trials were of three types. Stimulus S1 was presented by itself inside the RF, or S1 was presented along with S2 (both inside the RF), or S2 was presented by itself. S1, when present, was located at the estimated center of the RF (estimated online as the location eliciting the highest firing compared to baseline); S1 loom speed was fixed (6 °/sec). S2, when present, was positioned inside the RF such that its locations did not overlap with that of S1; S2 was chosen to be stronger than S1 (loom speed = 10 °/sec). Trials of the three types were interleaved randomly.

Stimuli were presented for 15 repetitions, with a duration of 250 ms each and an inter stimulus interval of 1.5 seconds.

### Data analysis and statistical tests

All analyses were done utilizing custom MATLAB scripts. Response firing rates were determined by counting the number of spikes over a time window following stimulus onset, and converting this count to firing rate (sp/s) after subtracting the baseline firing rate. For Imc, the window for computing firing rates started at 125 ms and ended at 275 ms, while for OT_10_ this window started at 100 ms and ended at 250 ms. These windows are similar or identical to those used in previous work in these areas ^13,19,20^. Average rates were calculated across all presentation repetitions after the removal of outliers, determined as values outside of the median ± 1.5*inter-quartile-range of the distribution. All statistical analyses were carried out within MATLAB. Parametric tests were used if data were normally distributed (tested using *lillietest*), non-parametric tests were used otherwise. Correction for multiple comparisons was performed using the Holm-Bonferroni method, if needed. Data shown as a ± b refer to mean ± s.e.m, unless specified otherwise. Data shown as CI = [a,b] refer to 95% confidence interval.

### Code and data availability

Software code and the data that support the findings of this study are available from the corresponding author upon reasonable request.

